# Genetic variation in parental effects contribute to the evolutionary potential of prey responses to predation risk

**DOI:** 10.1101/748251

**Authors:** Natasha Tigreros, Anurag A. Agrawal, Jennifer S. Thaler

## Abstract

Despite the ubiquity of parental effects and their potential impact on evolutionary dynamics, their contribution to the evolution of ecologically relevant adaptations remains poorly understood. Using quantitative genetics, here we demonstrate that parental effects contribute substantially to the evolutionary potential of larval antipredator responses in a leaf beetle (*Leptinotarsa decemlineata*). Previous research showed that larger *L. decemlineata* larvae elicit stronger antipredator responses, and mothers perceiving predators improved offspring responses by increasing intraclutch cannibalism –an extreme form of offspring provisioning. We now report substantial additive genetic variation underlying maternal ability to induce intraclutch cannibalism, indicating the potential of this adaptive maternal effect to evolve by natural selection. We also show that paternal size, a heritable trait, impacted larval responses to predation risk, but that larval responses themselves had little additive genetic variation. Together, these results demonstrate how larval responses to predation risk can evolve via two types of parental effects, both of which provide indirect sources of genetic variation for offspring traits.

## 1. INTRODUCTION

Parental effects, which are widespread in animals and plants, provide an important source of variation in offspring phenotype and fitness complementing that due to the direct inheritance of genes (Lande and Kirkpatrick 1990; Mousseau and Fox 1998; Wolf et al. 1998; Räsänen and Kruuk 2007). From the provisioning of parental care to the transfer of hormones and antibodies to young, parents alter the phenotype of their offspring, sometimes in an adaptive manner. Indeed, parental effects have been recognized as an important component of phenotypic variation that may facilitate rapid evolutionary responses to a number of ecological stressors (Mousseau and Fox 1998; Räsänen and Kruuk 2007; Donelson et al. 2018). Yet, predictions on the evolutionary consequences of parental effects are complicated by the fact that parental effects may be themselves shaped by the environmental conditions that parents experience, by genetic differences among parents, and by the interaction of these two (McAdam et al. 2014). While environmental and genetic influences in the parental generation should strongly impact evolutionary dynamics –increasing vs. decreasing trait response to selection–, little empirical work to date has partitioned their contribution to variation in offspring traits (reviewed in Räsänen & Kruuk 2007), especially those involving responses to natural ecological stressors.

Quantitative assessments of parental effects were initially included in quantitative genetic studies with the sole purpose of controlling for non-genetic sources of variation in offspring (Falconer and Mackay 1996; Lynch and Walsh 1998; Futuyma 2009). Nonetheless, it is now recognize that these effects often reflect genetic differences among parents, and therefore can evolve in response to selective forces occurring in both the parental and the offspring generation (Räsänen and Kruuk 2007). Importantly, genetic variation in parental effects provides an additional source of genetic variation that would facilitate evolution of offspring traits that hold little additive genetic variation (Räsänen and Kruuk 2007).

Parental effects on offspring often reflect environmental hardships that the parents experienced, including limited food, extreme weather, and a high risk of predation (Mousseau and Dingle 1991; Mousseau and Fox 1998). The triggering of environmental parental effects may reflect a passive consequence of stress or the resource environment that parents experience, or may involve adaptive responses counter to those conditions. Research over the last two decades indicates that parental effects can function as a form of adaptive transgenerational plasticity – commonly referred as “anticipatory parental effects” (Wade 1998; Agrawal et al. 1999; Galloway and Etterson 2007; Marshall and Uller 2007; Love and Williams 2008; Sheriff and Love 2013). Here, parents improve their offspring’s fitness by matching the offspring’s phenotype to environmental challenges they will likely experience (e.g. Marshall & Uller 2007; Sheriff & Love 2013). Despite growing evidence on the adaptive nature of parental effects on offspring (Agrawal et al. 1999; Sheriff et al. 2010; Storm and Lima 2010; Jensen et al. 2014), evidence of genetic variation in anticipatory parental effects (i.e. maternal genotype by environment interaction) is scarce in both animals (but see Fox et al. 1999) and plants (Galloway 2005). As a consequence, support for the evolutionary potential of anticipatory parental effects is to date limited (reviewed in Wade 1998; Räsänen and Kruuk 2007; McAdam et al. 2014).

Parental effects have been found to be important determinants of traits associated with predator avoidance in a number of species. For example, in a recent study in a leaf beetle (*Leptinotarsa decemlineata*), we demonstrated that variation in larval responses to predation risk was partially determined by an anticipatory parental effect. When experiencing predation risk, larger hatchlings were found to elicit stronger antipredator responses –measured as greater feeding reductions in the presence of predators. Remarkably, mothers increased offspring provisioning by inducing egg cannibalism within their clutches after detecting a high risk of predation. As a consequence, cannibalistic offspring –being larger and in better nutritional condition than their non-cannibal siblings– exhibited stronger responses to predation risk (Tigreros *et al.* 2017).

Here, we use classic quantitative genetics to examine the relative importance of parental effects for traits associated to predator avoidance in *L. decemlineata* larvae, including decreased foraging activity (leaf consumption) and increased assimilation efficiency. Specifically, we first estimate the contributions of maternal effects (V_M_), relative to that of additive genetic (V_A_) and environmental effects (V_E_), for larval responses to predation risk. Second, we test if larval responses to predation risk –which are known to depend on the larva’s initial size–, are influenced by variation in maternal or paternal body size, a key trait known to impact offspring phenotype in many systems (Fox 1994; Bernardo 1996*a*; Fox and Czesak 2000; Bennett and Murray 2014). Finally, we estimate the relative contribution of additive genetic and environmental variances (V_A_ and V_E_) of maternal responses to predation risk, including changes in intraclutch cannibalism –a mechanism linked to an anticipatory maternal effect.

## 2. METHODS

### (a) Breeding design

To estimate the relative contribution of parental effects to larval responses to predation risk we used a half-sib design (Falconer and Mackay 1996) (see Fig. 1). The experiment was initiated with 25 females collected from a field population in Ithaca, NY, which were allowed to lay eggs in the laboratory; their offspring, once mature, were considered the “parental generation” (Figure 1) from which sires and dams were selected. All sires, dams, and offspring from the different sire by dam crosses were reared separately from birth –which minimizes common environmental effects– and were maintained in standardized conditions, fed with *Solanum tuberosum* L (Yukon Gold variety) with 18-L: 6-D photoperiod and corresponding temperatures of 23: 21 °C. The half-sib families were initially established with 22 males (sires) each randomly mated to three unrelated females (dams). Sires that failed to inseminate at least two females were excluded and final analyses included fewer sires and dams, which is specified in each section below.

**Fig. 1.**
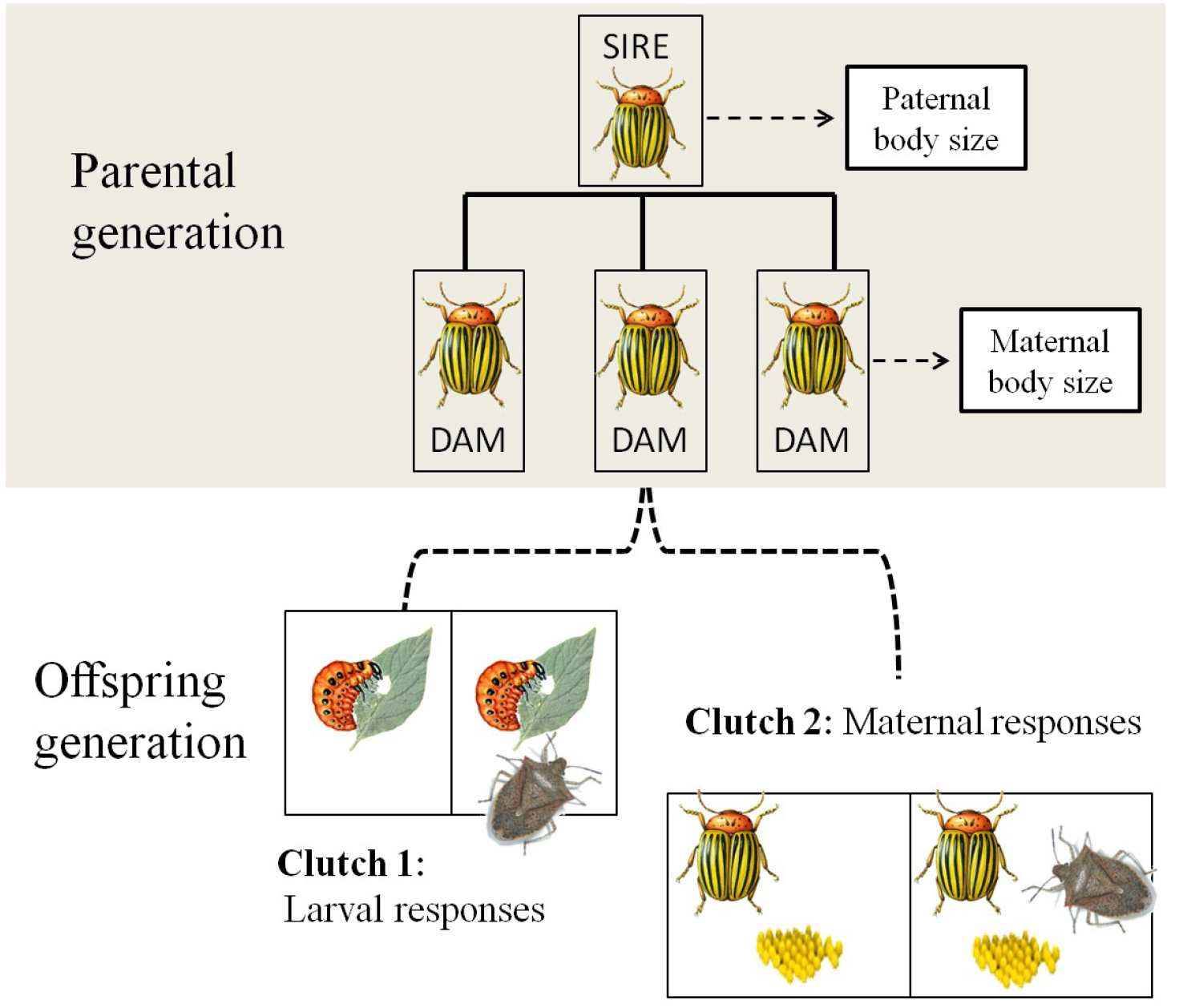
Diagram of *L. decemlineata* half-sib split-brood design showing one sire family. Nineteen sires were each mated to three virgin females (dams). The offspring of these three dams were used to investigate parental effects on plastic responses to predation risk: predator-free (P-Free) vs. predation risk (P-Risk). First, we examined the overall contribution of maternal effects (V_M_), relative to the additive genetic and environmental components (V_A_ and V_E_), to larval anti-predator responses: feeding reductions and increases in assimilation efficiency. Second, we compared how paternal and maternal body size influenced such larval responses to predation risk. Finally, using adult-mated-females, (from Clutch 2), we examined if there was additive genetic variation (V_A_) underpinning maternal response to predation risk. Maternal responses included changes in clutch size, proportion of viable offspring, and levels of intraclutch cannibalism; the last of these (increases in intraclutch cannibalism), is known to act as an adaptive maternal effect by improving larval anti-predator responses (Tigreros et al. 2017).

### (b) Maternal effects on larval responses to predation risk

In a first step, we followed a variance partitioning strategy to quantify the amount of variance in offspring’s traits –larval leaf consumption and assimilation efficiency– that is explained by maternal identity (V_M_), while accounting for the contributions of additive genetic inheritance (V_A_) and environmental variances (V_E_) (McAdam et al. 2014). Final analysis included 19 sires mated to 47 dams (with varying numbers of dams per sire). We estimated larval plastic responses to predation risk by measuring changes in larval feeding (leaf consumption) and assimilation efficiency in response to predation risk. To do this, six eggs from each maternal family (within the same clutch) were separated right before hatching and kept in individual 266 ml cups, which prevented intraclutch cannibalism and its influences on larval responses (Tigreros et al. 2017). Additionally, to control for potential effects due to hatching asynchrony within and among families, we recorded variation in egg pigment levels, Low, Medium, and High, as described in Tigreros et al. (2017).

Hatchlings from half of each maternal family were kept with a sham predator (“predation risk” environment), while the remaining siblings were kept without it, hence experiencing a “predator-free” environment (Fig. 1). Sham predators consisted of adult male *Podisus maculiventris* –a generalist stink bug that commonly feeds on *L. decemlineata* larvae– whose stylet’s terminal segment had been removed. While these altered stink bugs are no longer able to kill the beetle larvae, previous studies have shown that their behaviour and lifespan do not significantly differ from that of unaltered predators (Griffin and Thaler 2006; Thaler et al. 2012; Kaplan et al. 2014). Larvae were kept in the predator-free and predation risk environments for a total of three days (4-day old larvae). Then, we measured leaf consumption as consumed leaf area (mm^2^) using ImageJ software (version 1.45), and assimilation efficiency as the ratio of 4-d old larval mass (measured to the nearest 0.1mg using a Mettler AT261 balance; Mettler Toledo, Columbus, OH, USA) over amount of leaf consumed.

Statistical analysis, of maternal effects on larval responses to predation risk, involved the use of linear mixed models with maximum likelihood estimation. For each trait –leaf consumption and assimilation efficiency–, we included the treatment effect as a fixed factor (predator-free vs. predation risk) while the sire, dam, sire-by-treatment interaction and dam-by-treatment interaction (referred as GxE and MxE) were all included as random factors. Significance of variance components for larval responses to predation risk were estimated using the REML method (Proc MIXED) and likelihood ratio tests (Saxton and SAS Institute. 2004). Evidence of a significant GxE or MxE was investigated in more detail by testing the null hypothesis that genetic correlations of traits associated with plastic responses to predation risk, measured across the predator-free and predation risk environment (*r*_*A*_) were = 1 (Lynch and Walsh 1998; Messina and Fry 2003). Genetic correlations significantly less than 1 suggest the potential for independent trait evolution in different predation-risk environments and thus would provide additional support for GxE. Additionally, because expression of maternal and additive genetic effects is expected to differ within different environments, we estimated the genetic variance components (V_M_, V_A_, V_E_) and associated genetic parameters (e.g. narrow sense heritabilities and genetic coefficient of variation) for the larval traits within each environment –predator-free and predation risk (Table 1S). Variance components were calculated assuming the dominance variance to be zero: V_A_ = 4sire, V_M_ = dam – sire, and V_P_ = total phenotypic variance (Falconer and Mackay 1996). Narrow sense heritabilities (*h*^*2*^) of traits within a predator environment were then calculated as V_A_ / V_P_, representing the proportion of the total phenotypic variance explained by the direct genetic variance (Falconer and Mackay 1996).

**Table 1.**
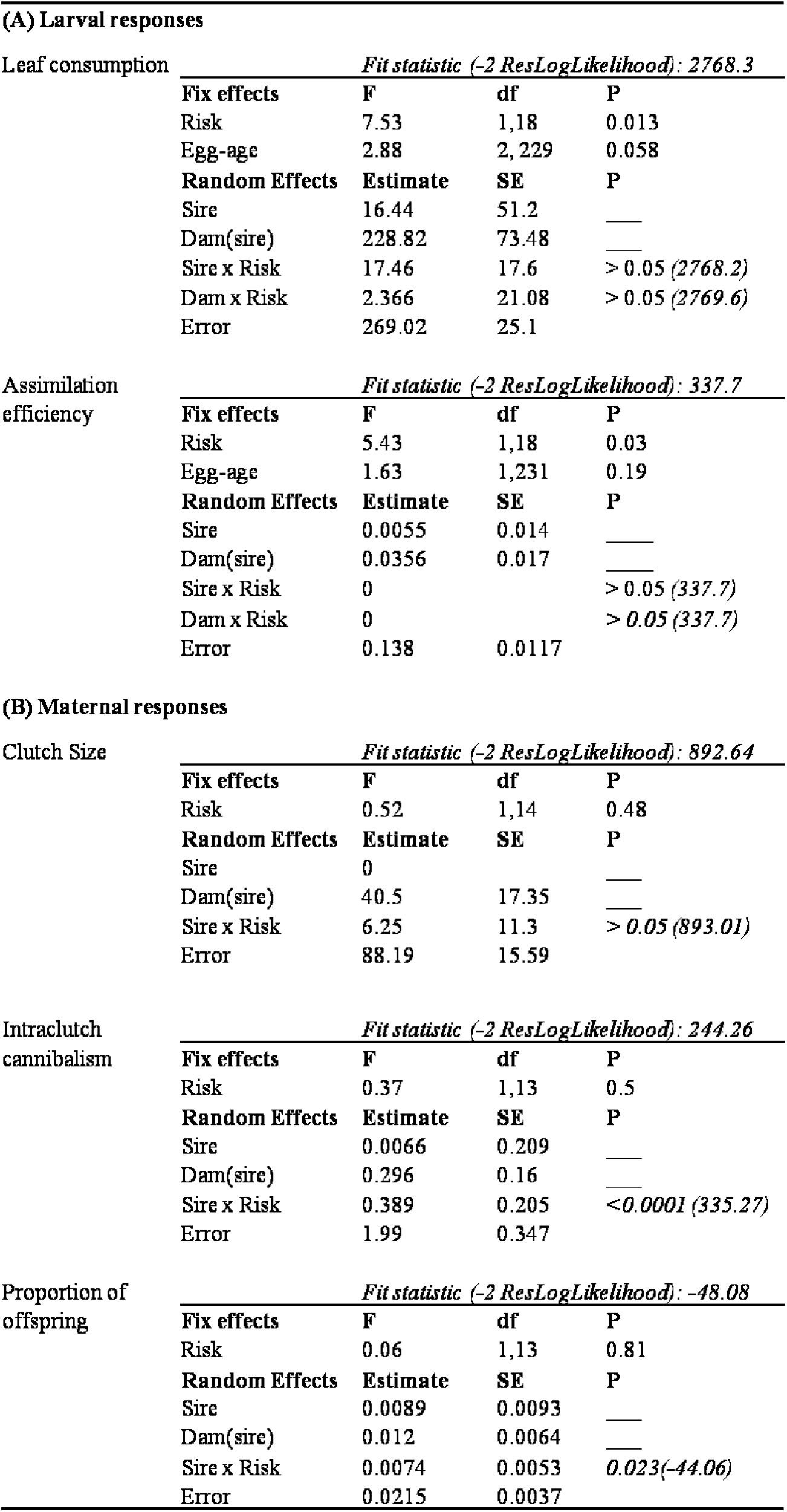
Tests on the statistical significance of the predation risk treatment (Risk) and its interaction with the dam (MxE) and sire (GxE) components. Interaction terms were tested using likelihood ratio test that compare the fit statistic,*-2 ResLog likelihood fit*, of the full model (showed on top row) with that of the model without the term of interest (showed in parenthesis after the P value).

### c) Parental size effects on larval responses to predation risk

In a second step, we followed a trait-based approach (McAdam et al. 2014) to investigate the effects of both maternal and paternal body size on larval plastic responses to predation risk. Here, the effects of specific parental traits, the body size of sires (fathers) and dams (mothers), are modeled. Because our main interest was to investigate effects on larval plastic responses to predation risk (rather than effects on the traits in each environment) we calculated larval plastic responses using 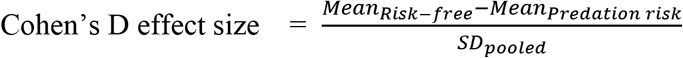 and used multiple regression analysis to test for both maternal and paternal size effects (for a similar approach see Bennett and Murray 2014). Given that previous studies have shown that larger hatchlings elicit stronger antipredator responses, we included larval size (averaged for each family) as an additional predictor. Additionally, we analyzed the effects of parental body size on larval responses using a “hybrid approach” (McAdam et al. 2014), which included sire and dam identities (as described in “maternal effects on larval responses to predation risk”) plus maternal (or paternal) body size and its interaction with predation risk treatment as fixed effects (Size_sire_ x E and Size_dam_ x E). This approach allow us to model the effects of specific parental traits (here body size) while accounting for the remaining variation in maternal and sire effects (V_M_ and V_A_). However, these more complex models, especially when unbalanced, may bias estimates of genetic parameters as well as inference for fixed effects (e.g. Kenward and Roger 1997; Stroup and Littell 2002).

### (d) Maternal plastic responses to predation risk

In a last experiment, we measured the amount of genetic variance (and associate genetic parameters) underlying maternal responses to predation risk including clutch size, levels of intraclutch cannibalism, and proportion of viable offspring within a clutch (Fig. 1). Note that although intraclutch cannibalism and clutch size may influence progeny phenotype (e.g. progeny size), these are here considered a maternal rather than an offspring trait (see Mousseau & Fox 1998). To measure maternal responses to predation risk, two females from each maternal family (and the same clutch) were individually reared to adults and mated with a full sibling. Because males in a number of insects can influence female reproduction (e.g. fecundity), mating full sibs may reduce chances to introduce an additional source of variation to the estimates of additive genetic variance in maternal traits. Importantly, mating full-sibs did not cause any apparent inbreeding effects in female reproduction: clutch size and offspring produced by full-sib pairs were comparable to those observed in females that had mated with unrelated males (see Inbreeding analysis in Supplemental information). After mating, half of the females were each kept with two sham predators (predation risk environment) in a 0.5 L cup, with abundant plant foliage for feeding and oviposition. The rest of females were kept in similar conditions but without the sham predators (predator-free environment). We collected the first two clutches that females laid and stored them individually with a fresh leaflet (in 30 ml cups). Clutch size (number of eggs) and offspring produced were measured in the same clutch while intraclutch cannibalism, the proportion of eggs that were fully consumed by the new hatchlings, were measured on a second clutch (Collie et al. 2013; Tigreros et al. 2017).

The statistical approach to examine maternal responses to predation risk was similar to that used for larval responses to predation risk but, because each dam included only one daughter per treatment, maternal effects on maternal responses to predation risk (V_M_ x E) cannot be estimated. Final analysis of maternal responses to predation risk included 15 sires mated to 32. Additionally, to estimate variance components for levels of intraclutch cannibalism we used a generalized linear model using LAPLACE method (Proc GLIMMIX), which captured the binomial distribution of intraclutch cannibalism using a logit link function (Saxton and SAS Institute. 2004). Because the residual variance estimates of this type of model is correlated with the mean of the population, we estimated heritability as “latent-scale heritability”, by adding the variance component related to the link function (Nakagawa and Schielzeth 2010; Calsbeek et al. 2015), which for binomial models is = 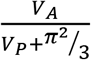. All analyses were conducted using SAS 9.4 (SAS Institute Inc., Cary, NC, USA).

## 3. RESULTS

### (a) Maternal effects on larval responses to predation risk

Larval plastic responses to predation risk were substantial, involving a 22% reduction in leaf consumption (F_1,18_ =7.53, P = 0.01; Fig. 2A, Table 1A) and a 17% increase in assimilation efficiency (F_1,18_ =5.43, P = 0.03; Fig. 2B, Table 1 A). Yet, we did not detect a significant interaction between predation risk and sire or dam families influencing the larval traits (Table 1A: Sire x Risk and Dam x Risk), which indicates that there was substantial plasticity in larval responses to predation risk, but there was not a detectable genetic or maternal variance underpinning larval responses.

**Figure 2.**
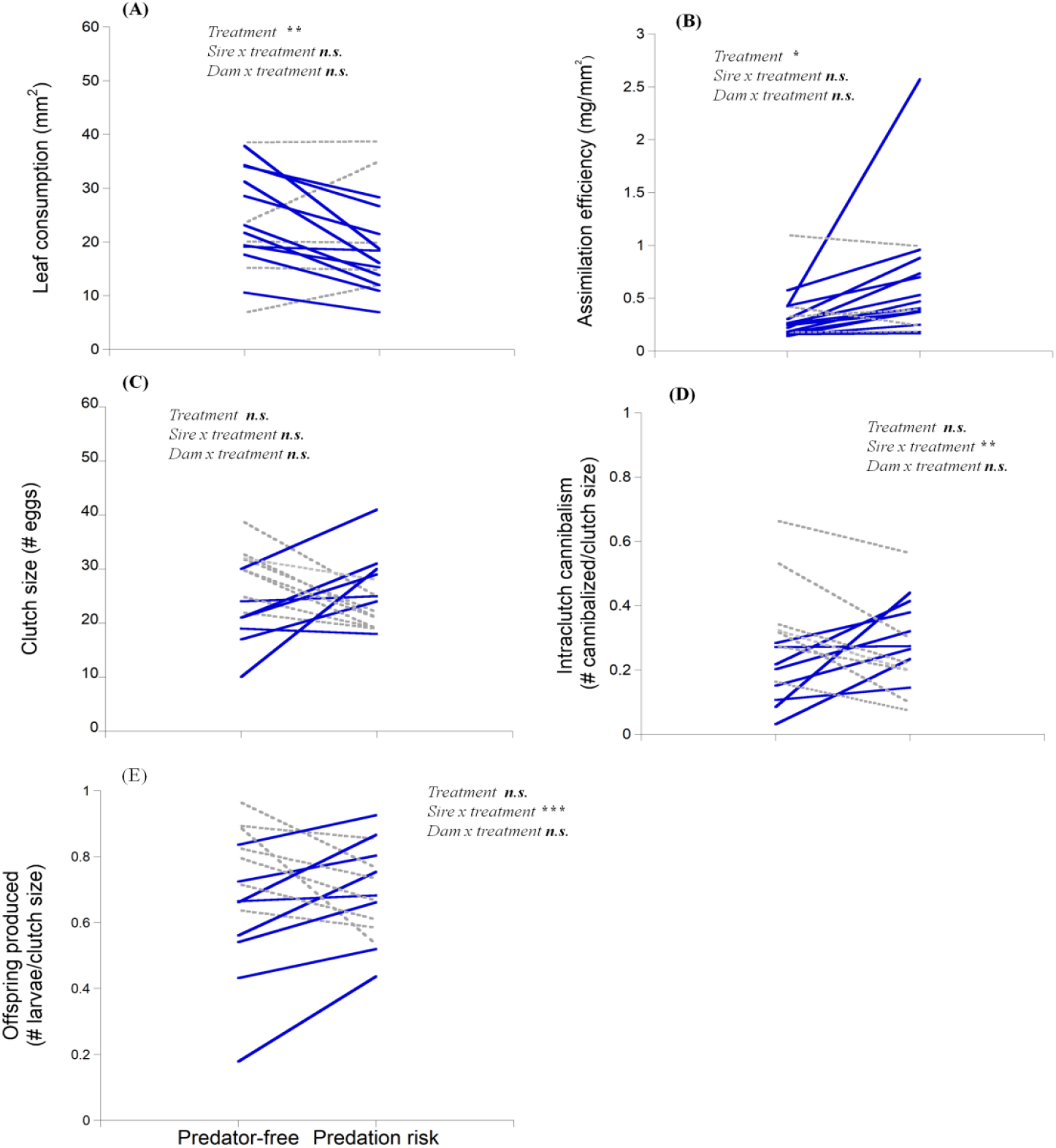
Reaction norms for *L. decemlineata* describing plastic responses to predation risk in terms of (A) Larval leaf consumption, (B) Larval assimilation efficiency, (C) Adult clutch size, (D) Intraclutch cannibalism and (E) Offspring number per clutch. Each line represents the mean score for each sire family. The slope of the line is a graphical representation of the strength and direction of plastic responses to predation risk shown by that family. Highlighted with solid blue lines (vs. doted grey lines) are the families that responded to predation risk in a direction that would typically improve offspring fitness (see Table 1 for further statistical elaboration).

Partitioning of variance components of larval traits within each predator environment (predator-free and predation risk environment) showed, as expected, differences in the relative importance of maternal (V_M_) and additive genetic effects (V_A_) (Fig. 3; Table 1S). While maternal effects (V_M_) explained a large proportion of the phenotypic variance in both environments, this was even stronger when in the predation risk environment, explaining over 50% of the variation in leaf consumption and about 30% in assimilation efficiency (Fig. 3; Table S1 in Supplemental information). In contrast, levels of additive genetic variation (V_A_) were not statistically significant in either environment (Table S1). Note, however, that the number of sire families in our study was close to the minimum recommended to detect V_A_ (Conner and Hartl 2004) and therefore, finding no additive genetic variance in larval traits could be due to a low statistical power rather than an absolute lack of genetic variance.

**Fig. 3.**
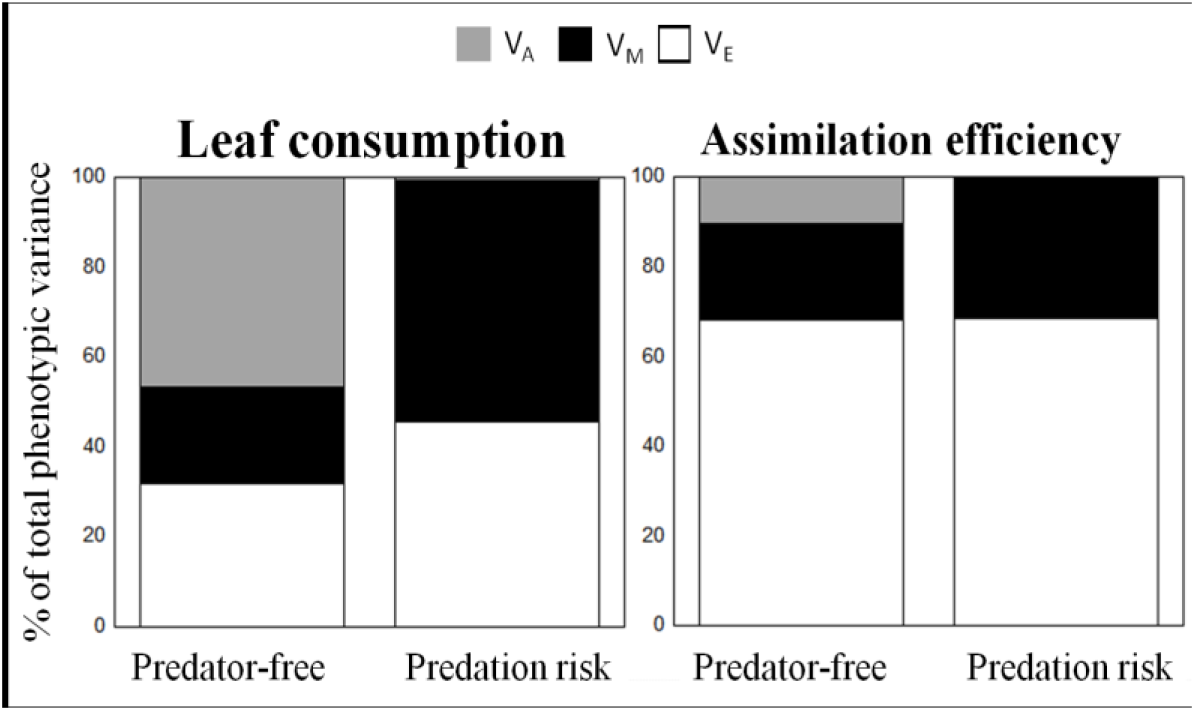
Percent of phenotypic variation (for leaf consumption and assimilation efficiency) explained by the different genetic components of variation within each predator. Genetic components of variation include: additive genetic (V_A_), maternal (M_A_), and environmental (V_E_) effects.

### (b) Parental size effects on larval responses to predation risk

To test parental size effects on larval responses, we first partitioned variation of adult body size into additive genetic and environmental components. These results revealed a significant additive genetic component (V_A_ = 0.078 ± 0.056, *p=0.025*), with moderate heritability levels (*h*^*2*^ = 0.2), underlying body size in adults.

Evaluation of parental size effects on larval responses provided similar results using both the trait-based approach of regressing parents’ body size on larval responses– and the hybrid approach of including parent’s body size as covariate plus Sire and Dam identities (McAdam et al. 2014). Regression analyses (controlling offspring size), revealed that paternal body size, but not maternal body size, influenced both the magnitude and direction of larval responses to predation risk including leaf consumption (*R*^*2*^ =0.26: ßsire= −0.51, *p*= 0.001; ßdam= 0.04, *p*= 0.25; ßlarvae= 0.06, *p*= 0.4; Fig. 4A), and assimilation efficiency (*R*^*2*^ =0.23: ßsire= 0.48, *p*= 0.003; ßdam=-0.05; *p*= 0.73; ßlarvae=0.04, *p*= 0.77; Fig. 4B). Smaller sires producing larvae with the strongest responses (reduced feeding and increased assimilation efficiency). Results from models that included sire and dam identity corroborated the effect of sire size on larval responses (Table S2 in Supplemental information).

**Fig. 4.**
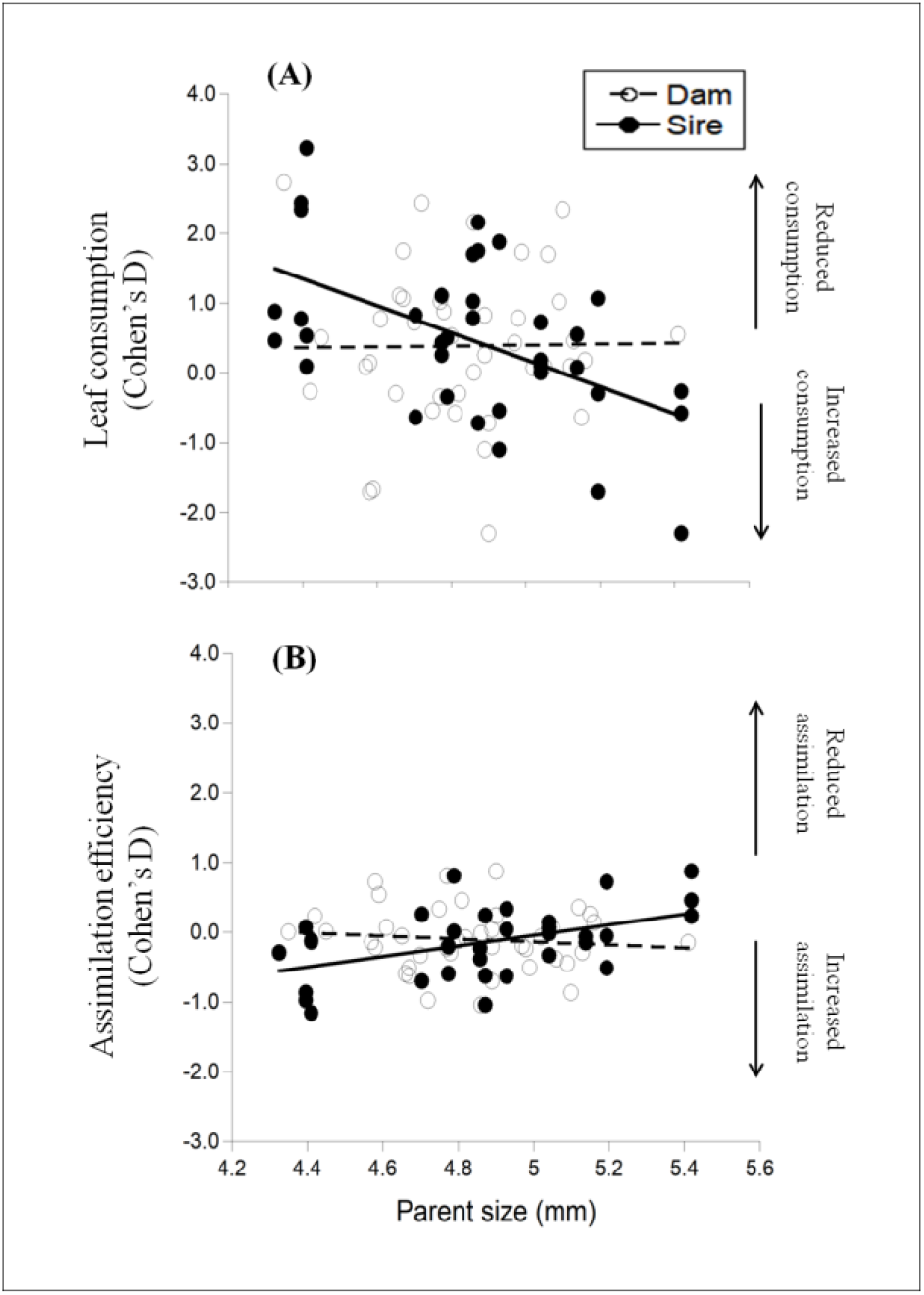
Parental body size effects on larval responses to predation risk for (A) leaf consumption and (B) assimilation efficiency in *L. decemlineata* larvae. The magnitude and direction of larval plastic responses were calculated using Cohen’s D effect size 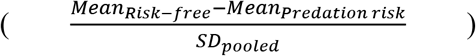. Each observation represents the effect size and mean body size for each maternal family (full siblings).

### (b) Maternal response to predation risk

Analyses of maternal responses to predation risk indicated that there was not a main effect of predation risk treatment on clutch size, intraclutch cannibalism, or number of viable offspring produced per clutch (Table 1B; Fig. 2C–E). However, there was a significant sire by predation treatment interaction for proportion of intraclutch cannibalism and number of viable larvae produced, indicating that there is substantial genetic variation for these maternal traits, involving differences in the magnitude as well the direction of the response (Fig. 2 D, E). Further investigation of this GxE, by testing significance of genetic correlations across the predator-free and predation risk environments, indicated that these were significantly less than 1 (intraclutch cannibalism *r*_*A*_ = −0.01, se = 0.7, *p* = 0.03; proportion of offspring *r*_*A*_ = 0.74, se = 0.5, *p* < 0.0001). These results suggest the potential for independent trait evolution in the different predation environments, providing additional support for GxE in maternal responses to predation risk (Lynch and Walsh 1998).

## DISCUSSION

Parental effects are important determinants of traits associated with predator avoidance in a number of species. As for all phenotypes, parental effects are shaped by the environment, the genotype, and their interaction. Only by disentangling the relative contribution of such different sources of parental effects –e.g., those owed to the environment vs. the parent’s genes– we can understand their role in organisms’ evolution.

As observed in previous studies of *L. decemlineata* (Hermann and Thaler 2014; Kaplan et al. 2014; Tigreros et al. 2017, 2018), we found strong plastic responses to predation risk, involving 28% reductions in leaf consumption and 15% increases in assimilation efficiency. Coupling of feeding reductions with increased assimilation efficiency is critical to prey fitness, as this allows prey to lower chances of predation while minimizing costs associated with reduced-food intake (Thaler et al. 2012; Kaplan et al. 2014). Based on 19 sires, we found little support of an additive genetic or a maternal component underpinning larval responses to predation risk. However, analyses that included size of the parents as covariates revealed that variation in responses –for both leaf consumption and assimilation efficiency– were at least partially explained by paternal size. Specifically, smaller fathers produced larvae with the greatest plasticity in response to predation risk –strong feeding reductions and increases in assimilation efficiency. Given that females often provide more reproductive investment than males, studies on parental effects often rely on the notion that the maternal phenotype is the main influence on the offspring. However, as we found in this study, male body size can be linked to parental performance (Eilertsen et al. 2009), perhaps through changes in the quality of their ejaculates (Gillott 2003), which is known to include accessory glands compounds in *L. decemlineata* beetles (Loof and Lagasse 1972). Independent of the exact mechanism, because adult body size had a significant additive genetic component, size related paternal effects represented an indirect source of genetic variation shaping larval plastic responses to predation risk.

Estimates of maternal variance (V_M_) explained about half of the phenotypic variance in leaf consumption when expressed under high risk of predation. These results are concordant with the notion that the contribution of parental effects, relative to the environmental and additive genetic variance components, is high for traits that are expressed early in life (Bernardo 1996*b*; Lindholm et al. 2006; Wilson and Reale 2006; White and Wilson 2019) and under environmental stress (e.g. Rudin-Bitterli, Mitchell & Evans 2018).

In a previous study we demonstrated that larval responses to predation risk (including decreases in leaf consumption) was improved through an anticipatory parental effect: mothers experiencing the risk of predation increased offspring provisioning by inducing intraclutch cannibalism (Tigreros et al. 2017); cannibals, in better nutritional condition than non-cannibal siblings, exhibited stronger antipredator behaviors. Here, we found that mothers indeed responded to predation risk by altering levels of egg cannibalism within their clutches. However, such responses varied in magnitude and direction, with some families increasing and others decreasing cannibalism. In *L. decemlineata*, intraclutch cannibalism results in a classic life history tradeoff between investment in offspring quality (cannibalistic offspring) and quantity (cannibalized offspring). While fitness of the individual offspring may always improve with cannibalism, optimal levels of cannibalism within a clutch should reflect the number of cannibalistic offspring that would maximize female reproductive success in a given environment (e.g. under predation risk). Finding that families with the highest and lowest levels of intraclutch cannibalism –in the predator-free environment– responded to predation risk by decreasing and increasing cannibalism respectively, suggests that intermediate levels of intraclutch cannibalism may be optimal under environments with high predation risk. Importantly, such variation in maternal responses to predation risk (changes in intraclutch cannibalism and offspring produced) reflected genetic differences among the mothers (significant GxE) indicating the potential for evolutionary change of a maternal effect, in response to predator.

The conditions under which organisms rely on parental effects vs. within-generation phenotypic plasticity remains an open question in evolutionary ecology. Using plasticity alone, *L. decemlineata* larvae can achieve substantial feeding reductions (e.g. ~28% in this study) when facing predation risk. However, our previous work indicates that these responses are constrained by the larvae’s nutritional state (Tigreros et al. 2017), and larvae feeding on lower quality host plants have shown weaker feeding reductions in response to predation risk (Kaplan et al. 2014; Tigreros et al. 2017). Through cannibalism, *L. decemlineata* appears to overcome such nutritional constraints, and cannibals that experience predation risk are capable of reducing leaf consumption, even on low quality host plants. Thus, this extreme type of maternal investment should improve offspring fitness when facing highly stressful environments (Marshall et al. 2008; Olofsson et al. 2009).

Recent theoretical evidence suggests that evolution of parental effects is promoted when there is strong selection on the phenotype and when within-generation plasticity is constrained (Auld et al. 2010; Kuijper and Hoyle 2015). Predation is unquestionably one of the strongest selective forces in nature, and is therefore thus likely to deplete genetic variation in traits associated with predator avoidance. In contrast, genetic variation underlying paternal and maternal effects is expected to remain “protected” from the eroding effects of selection when carried by the other sex (Wade 1998). Accordingly, results from this study indicate that even if larval responses to predation risk were holding little or no genetic variation, parental effects –via induced intraclutch cannibalism and paternal body size– should provide the necessary genetic variation for future natural selection to act upon (Räsänen and Kruuk 2007). Additionally, because *L. decemlineata* larval nutritional condition is a function of host plant quality as well as the degree of maternal provisioning, evolution of paternal effects may be favored in environments where larval plasticity is constrained by poor host plant quality. Together, our results show that the evolutionary potential of predator avoidance in *L. decemlineata*, relies on at least two different genetic parental effects, one linked to paternal body size and the other to maternal induction of intraclutch cannibalism.

## ACKNOWLEDGEMENTS

We thank Rachel H. Norris for help with collection of data. We also thank Kyle Benowitz and Nicholas Aflitto, and two anonymous reviewers for their valuable comments on the manuscript. This work was supported by USDA grants to JST: NIFA 2013-02649, Federal Capacity Funds 1397484; and AFRI 2018-67013-28068.

