## Supplemental information 1_ for "Genetic variation in parental effects contribute to the evolutionary potential of prey responses to predation risk"

**Supplementary information 1.**

COMPARING OUTBRED (n1) WITH INBRED (N2) INDIVIDUALS FROM THE SAME POPULATIONS AND SAME YEAR.

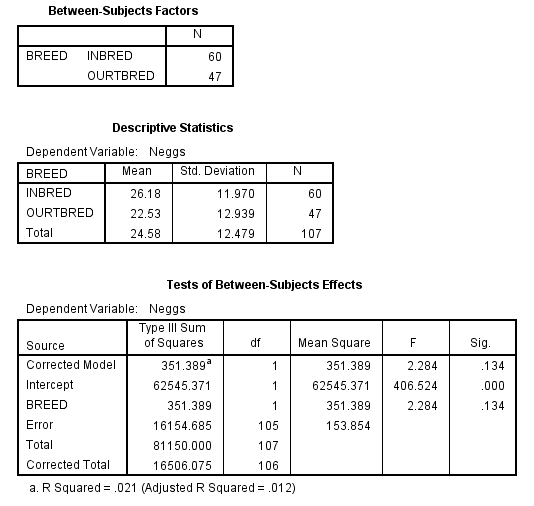

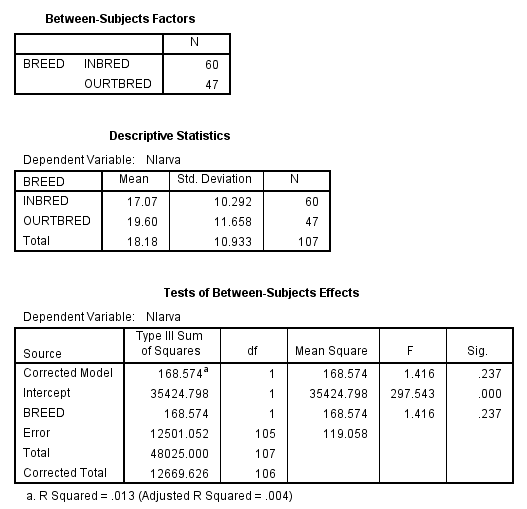
