## Supplemental Data 1 for "Genetic variation in parental effects contribute to the evolutionary potential of prey responses to predation risk"

**Table S2.** We estimated maternal size effects on larval responses using a “hybrid approach”, as described in (McAdam, Garant and Wilson, 2014). Here, the effects of a maternal trait on the offspring (in our case, the interaction of Dam’s body size with treatment) are modeled together with the maternal by treatment interaction (Dam x Risk)_._ This last term would capture the remaining “unknown maternal effects”. Similarly, we modeled paternal size effects on larval responses by including the interaction of treatment effects with sire’s body size as well as the interaction with sire identity.

|  |  | **Leaf consumption** | | |  | **Assimilation Efficiency** | | |
| --- | --- | --- | --- | --- | --- | --- | --- | --- |
| ***Dam size x Risk*** | | *Fit statistic (-2 ResLogLikelihood): 2586.7* | | |  | *Fit statistic (-2 ResLogLikelihood): 320.5* | | |
|  | **Fix effects** | **F** | **df** | **P** |  | **F** | **df** | **P** |
|  | Risk | 0.32 | 1,17 | 0.8 |  | 0.14 | 1,17 | 0.71 |
|  | Dam size | 223 | 1,215 | 0.14 |  | 0.09 | 1,217 | 0.77 |
|  | Dam size x Risk | 0.19 | 1,215 | 0.66 |  | 0.22 | 1,217 | 0.64 |
|  | Egg-age | 3.27 | 2,215 | 0.04 |  | 1.92 | 2 | 0.15 |
|  | **Random Effects** | **Estimate** | **SE** |  |  | **Estimate** | **SE** |  |
|  | Sire | 14.925 | 63.37 |  |  | 0.006 | 0.016 |  |
|  | Dam(sire) | 227.37 | 80.66 |  |  | 0.035 | 0.018 |  |
|  | Sire x Risk | 28.66 | 21.05 |  |  | 0.0008 | 0.0059 |  |
|  | Dam x Risk | 0 |  |  |  | 0 |  |  |
|  | Error | 276.14 | 25.25 |  |  | 0.14 | 0.013 |  |
| ***Sire size x Risk*** | | *Fit statistic (-2 ResLogLikelihood): 2477.8* | | |  | *Fit statistic (-2 ResLogLikelihood): 288.5* | | |
|  | **Fix effects** | **F** | **df** | **P** |  | **F** | **df** | **P** |
|  | Risk | 5.16 | 1,13 | 0.04 |  | 4.38 | 1,13 | 0.056 |
|  | Sire size | 0.47 | 1,205 | 0.49 |  | 0.07 | 1,207 | 0.797 |
|  | Sire size x Risk | 4.59 | 1,205 | 0.03 |  | 3.97 | 1,207 | 0.047 |
|  | Egg-age | 2.07 | 2,205 | 0.1 |  | 2.09 | 2,207 | 0.126 |
|  | **Random Effects** | **Estimate** | **SE** |  |  | **Estimate** | **SE** |  |
|  | Sire | 25.11 | 63.48 |  |  | 0.00815 | 0.015 |  |
|  | Dam(sire) | 247.34 | 84.71 |  |  | 0.035 | 0.016 |  |
|  | Sire x Risk | 10.431 | 17.3 |  |  | 0 |  |  |
|  | Dam x Risk | 1.627 | 23.28 |  |  | 0 |  |  |
|  | Error | 285.78 | 28.14 |  |  | 0.13 | 0.012 |  |
