## Supplemental Data 2 for "Genetic variation in parental effects contribute to the evolutionary potential of prey responses to predation risk"

**Table S1.** Showing genetic components of variation and associated genetic parameters for larval traits, within a predator environment: predator-free vs. predation risk.

| **(A) Larval traits** |  |  | |  |  | |  |  | |  | | |  | |  | |  |
| --- | --- | --- | --- | --- | --- | --- | --- | --- | --- | --- | --- | --- | --- | --- | --- | --- | --- |
|  |  | **Observational** | | | **Genetic** | | | **SE** | **p** | | **% of Vp** | | | | | |  |
| **Leaf intake** | Predator-free | V_Sire_ = | 69.27 | | V_A_ = | 277.06 | | 274.56 | 0.13 | | 46.78 | | | **Trait mean =** | | 29.47 | |
|  |  | V_Dam_ = | 195.18 | | V_M_ = | 125.91 | | 127.06 | 0.16 | | 21.26 | | | **CV% =** | | 56.48 | |
|  |  | V_E_ = | 327.86 | | V_E_ = | 189.33 | | 144.43 |  | | 31.96 | | | ***h² =*** | | 0.16 | |
|  |  | V_P_ = | 592.31 | |  |  | |  |  | |  | | |  | | 0.47 | |
|  | Predation Risk | V_Sire_ = | 0.62 | | V_A_ = | 2.48 | | 193.52 | n.s. | | 0.53 | | | **Trait mean =** | | 22.94 | |
|  |  | V_Dam_ = | 254.24 | | V_M_ = | 253.62 | | 117.56 | 0.009 | | 53.89 | | | **CV% =** | | 6.87 | |
|  |  | V_E_ = | 215.49 | | V_E_ = | 214.25 | | 100.70 |  | | 45.52 | | | ***h² =*** | | 0.00 | |
|  |  | V_P_ = | 470.65 | |  |  | |  |  | |  | | | |  | |  |
| **Assimilation** | Predator-free | V_Sire_ = | | 0.00 | V_A_ = | 0.01 | | 876.76 | n.s. | | 10.53 | | | **Trait mean =** | | | 0.66 |
| **efficiency** |  | V_Dam_ = | | 0.03 | V_M_ = | 0.03 | | 455.43 | 0.12 | | 21.46 | | | **CV% =** | | | 17.67 |
|  |  | V_E_ = | | 0.09 | V_E_ = | 0.09 | | 528.04 |  | | 68.01 | | | ***h² =*** | | | 0.09 |
|  |  | V_P_ = | | 0.13 |  |  | |  |  | | |  | |  | | | 0.11 |
|  | Predation Risk | V_Sire_ = | | -0.01 | V_A_ = | -0.04 | | 0.28 | n.s. | | 0.00 | | | **Trait mean =** | | | 0.58 |
|  |  | V_Dam_ = | | 0.06 | V_M_ = | 0.06 | | 0.19 | 0.01 | | 31.45 | | | **CV% =** | | | 0.00 |
|  |  | V_E_ = | | 0.13 | V_E_ = | 0.13 | | 0.19 |  | | 68.55 | | | ***h² =*** | | | 0.00 |
|  |  | V_P_ = | | 0.19 |  | |  |  | |  | | |  | |  | |  |
